# Aberration correction in STED microscopy to increase imaging depth in living brain tissue

**DOI:** 10.1101/2021.01.05.425408

**Authors:** Stéphane Bancelin, Luc Mercier, Emanuele Murana, Valentin Nägerl

## Abstract

We demonstrate an approach based on adaptive optics to improve the spatial resolution of STED microscopy inside thick biological tissue by a priori correction of spherical aberrations as a function of imaging depth. We first measured the aberrations in a phantom sample of gold and fluorescent nanoparticles suspended in an agarose gel with a refractive index closely matching living brain tissue. Using a spatial light modulator to apply corrective phase shifts, we imaged neurons in living brain slices and show that the corrections can substantially increase image quality. Specifically, we could measure structures as small as 80 nm at a depth of 90 *μ*m inside the biological tissue, and obtain a 60% signal increase after correction.

## Introduction

The advent of super-resolution microscopy, breaking the diffraction barrier of optical microscopy, has opened up tremendous opportunities for nanoscale imaging of live cells [1,2]. In particular, Stimulated Emission Depletion (STED) microscopy has made it possible to reconcile nanoscale resolution with imaging in live brain tissue preparations, setting new standards for visualizing the dynamic morphology of neurons [3–5].

STED microscopy relies on the use of two co-propagating laser beams: a Gaussian beam to excite fluorescent molecules within a diffraction-limited focal spot, and a depletion beam to de-excite the fluorescence of the peripheral molecules via stimulated emission. This joint action increases the spatial resolution by up to an order of magnitude [6]. Using appropriate beam shaping, STED microscopy can be operated either in 2D or 3D mode, typically using so-called donut or bottle beams. Applied in combination, these two modes allow for optimizing lateral as well as axial resolution at the same time [7–9].

STED microscopy depends crucially on the quality of the point-spread function (PSF) of the depletion beam. However, maintaining the STED PSF deep inside thick brain tissue preparations remains challenging. Indeed, optical aberrations, stemming from the optics and the specimen, induce distortions on the laser wavefront that can severely degrade the quality of the STED PSF (especially its symmetry and the null in the centre) [10–12]. While system aberrations tend to be static and can be compensated using appropriate optics, the sample-induced aberrations are more problematic and usually limit the ability to image much beyond the surface of the sample. However, in the last few years, several studies have addressed this challenge in various ways, e.g. combining STED with 2P excitation [13,14], objective lenses with correction collar [4], Bessel beams [15], optical clearing/index matching [16] or adaptive optics (AO) [11].

The use of AO, which makes it possible to pre-distort the wavefront and thus cancel both system and sample aberrations [17], is a promising way to preserve a high-quality PSF deep inside scattering tissue [18]. Spatial light modulators (SLM) offer a versatile solution not only to generate the focal STED donut [19,20], but also to implement aberration correction measures [7,8,11,21,22]. However, most of these techniques operate in closed loop [11], which necessitates numerous iterations to achieve appreciable correction. This imposes long exposure times to laser light on the sample, inducing a significant amount of photo-bleaching and toxicity. While an iterative approach is possible in settings where bleaching is very low or absent, as in the case of super-resolution shadow imaging [23], it is impractical when imaging positively labelled structures, which tend to bleach rapidly.

In this work, we propose a simple and robust approach to improve resolution in depth by using an a priori estimation of aberrations as a function of imaging depth, focusing on spherical aberrations, which are the main type of aberration induced by biological specimen. To that end, we measured the aberrations as a function of depth in an agarose gel containing gold and fluorescent beads and with a refractive index closely adjusted to the one of living brain tissue. We then show that these calibrated values can be used to pre-program an SLM at specified depths to correct the aberrations and to improve image quality, facilitating the depth penetration for nanoscale imaging of neuronal morphology in living brain tissue.

## Material and Methods

### STED microscope

Imaging was performed using a custom-built upright STED microscope based on pulsed excitation at 900 nm and pulsed depletion at 592 nm (Fig. 1a). Briefly, two-photon excitation (2P) is achieved using a femtosecond mode-locked Titanium:Sapphire laser (Tsunami, Spectra Physics) delivering 100 fs pulses at a 80 MHz repetition rate. Laser power was adjusted using a Pockels cell (302 RM, Conoptics). The STED beam was provided by a pulsed laser (Katana 06 HP, NKT Photonics) delivering 400 ps pulses at 80 MHz. Laser power was adjusted using a combination of a half-wave plate and a beam splitting polarizer. Both lasers were synchronized using ‘lock-to-clock’ electronics (Model 3930 and 3931, Spectra Physics).

**Fig. 1.**
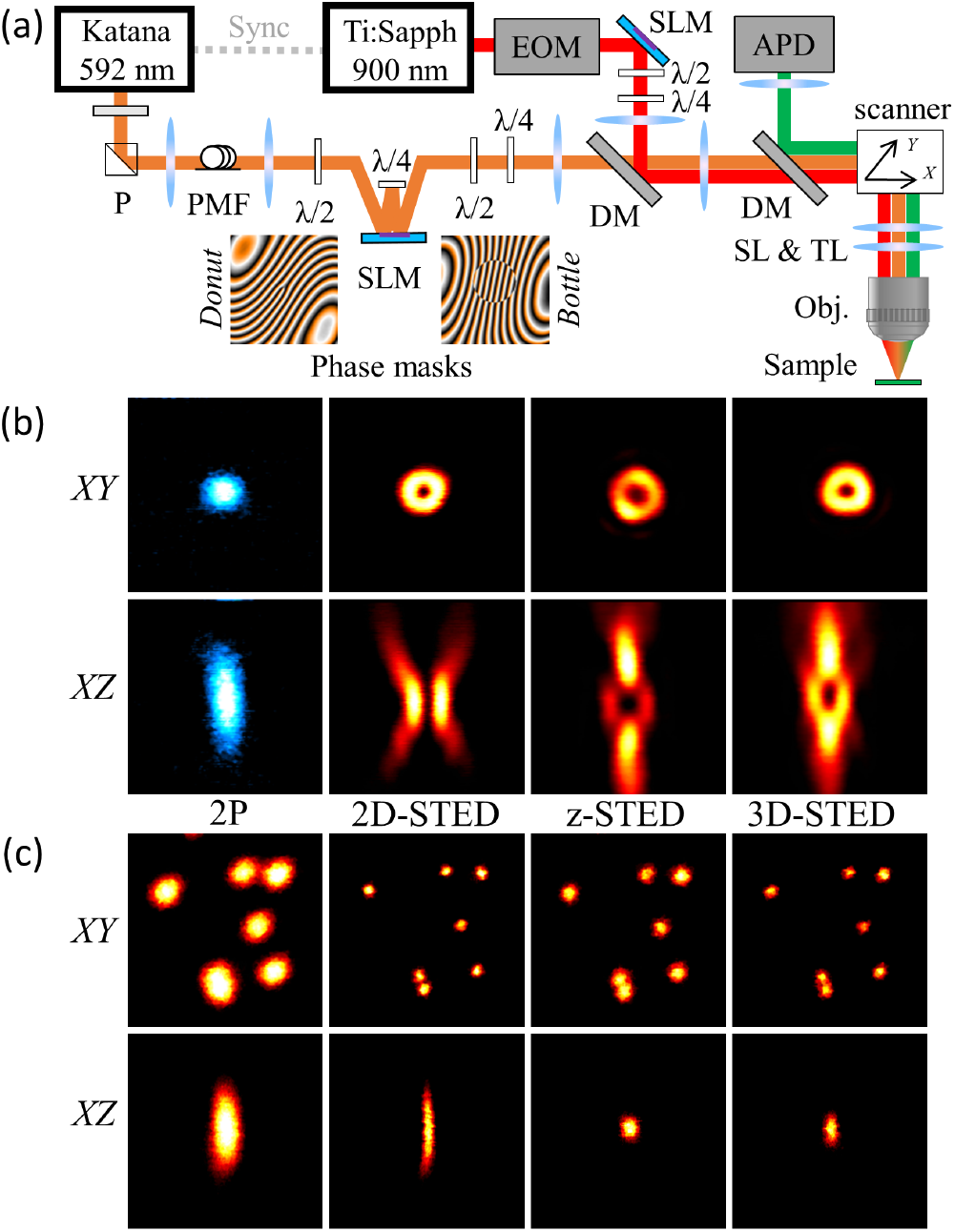
a) Schematic of the setup used for 2P-STED microscopy. P: polarizer, PMF: polarization maintaining fiber, SLM: spatial light modulator, *λ*/2 and *λ*/4: half and quarter wave-plates, DM: dichroic mirror, SL and TL: tube and scan lens, EOM: electro-optic modulator, APD: avalanche photodiode. b) 2P and STED beam PSFs obtained by imaging gold beads. c) Lateral and axial view of the effective PSFs assessed on 170-nm fluorescent beads. Image size: 3 × 3μ*m*^2^.

The STED beam was spectrally cleaned up using a band-pass filter (593/40, Semrock) and appropriately shaped using a ‘3D module’ (Aberrior Instruments), controlled through Imspector software, based on an SLM to generate a collinear mix of donut and bottle beams following the scheme described in Lenz *et al*. [7]. In this configuration, the SLM served two purposes: shaping the donut and bottle beams and correcting for optical aberrations. Half and quarter-wave plates (λ/2 and λ/4) were used to adjust the polarization to left-handed circular. The 2P and STED laser beams were combined using a long-pass dichroic mirror (DCSPXRUV –T700, AHF). Appropriate lens combinations were used to conjugate the SLM on a telecentric scanner (Yanus IV, TILL Photonics), which projected both scan axes on the back focal plane (BFP) of the objective lens (UPLanSAPO, 60x, Silicone Oil immersion, NA 1.3 Olympus) mounted on a z-focusing piezo actuator (Pifoc 725.2CD, Physik Instrumente). The epi-fluorescence signal was descanned, separated from incident beams using a long-pass dichroic mirror (580 DCXRUV, AHF) and detected by an avalanche photodiode (SPCM-AQRH-14-FC, Excelitas) with appropriate notch (594S-25, Semrock) and bandpass filters (680SP-25, 520-50, Semrock) along the emission path. Signal detection and hardware control were performed with the Imspector scanning software (Abberior Instruments) via a data acquisition card (PCIe-6259, National Instruments).

To assess donut or bottle beam quality, a pellicle beam splitter (BP145B1, Thorlabs) was flipped into the beam path to detect the signal reflected on gold beads (150 nm Gold nanospheres, Sigma Aldrich) on a photomultiplier tube (MD963, Excelitas). The STED profile was changed from donut to bottle by rotating the half-wave plate placed before the SLM. In the following, 2D-STED, z-STED and 3D-STED will refer to images acquired using a pure donut, pure bottle or a combination of donut and bottle beams, respectively (Fig. 1b). Optical resolution was assessed by imaging fluorescent beads (yellow-green fluorescent beads, 40 nm or 170 nm in diameter, Invitrogen) immobilized on glass slides (Fig. 1c)

### STED beam quality and aberration correction

In the context of thick biological samples imaging, optical aberrations are defined as the deviation of the wavefront from its perfect shape, which is commonly expressed as a series of Zernike polynomials:

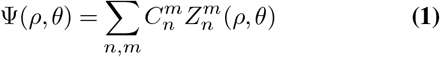

where Ψ is the aberration function, 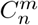 and 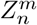 are the Zernike coefficients and polynomials respectively (Table 1). Zernike polynomials are widely used to model aberrations because they are orthogonal and each polynomial function corresponds to an optical aberration, such as astigmatism, coma, or spherical, in the paraxial approximation [11,24].

**Table 1.** First orders Zernike polynomials

| n | m | Formula | Aberration type |
| --- | --- | --- | --- |
| 0 | 0 | 1 | Piston |
| 1 | 1 | $2\rho\cos(\theta)$ | Tip |
| 1 | -1 | $2\rho\sin(\theta)$ | Tilt |
| 2 | 0 | $\sqrt{3}(2\rho^2 - 1)$ | Defocus |
| 2 | 2 | $\sqrt{6}\rho^2\cos(2\theta)$ | Astigmatism |
| 2 | -2 | $\sqrt{6}\rho^2\sin(2\theta)$ | Astigmatism |
| 3 | 1 | $2\sqrt{2}(3\rho^3 - 2\rho)\cos(\theta)$ | Coma |
| 3 | -1 | $2\sqrt{2}(3\rho^3 - 2\rho)\sin(\theta)$ | Coma |
| 3 | 3 | $2\sqrt{2}\rho^3\cos(3\theta)$ | Trefoil |
| 3 | -3 | $2\sqrt{2}\rho^3\sin(3\theta)$ | Trefoil |
| 4 | 0 | $\sqrt{5}(6\rho^4 - 6\rho^2 + 1)$ | Spherical |

Fig. 2 shows the effect of these first-order aberrations on the STED beam profile. N otably, e ach a berration presents a specific e ffect t hat c an b e e asily i dentified by lo oking at the STED beam distortion. Importantly, when using a high NA objective, as in STED microscopy, the paraxial approximation is not valid anymore, which leads to an appreciable coupling between the modes. For example, in Fig. 2, the spherical aberration is accompanied by a shift in the center of the donut beam, which reflects the effect of classical tip 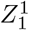.

**Fig. 2.**
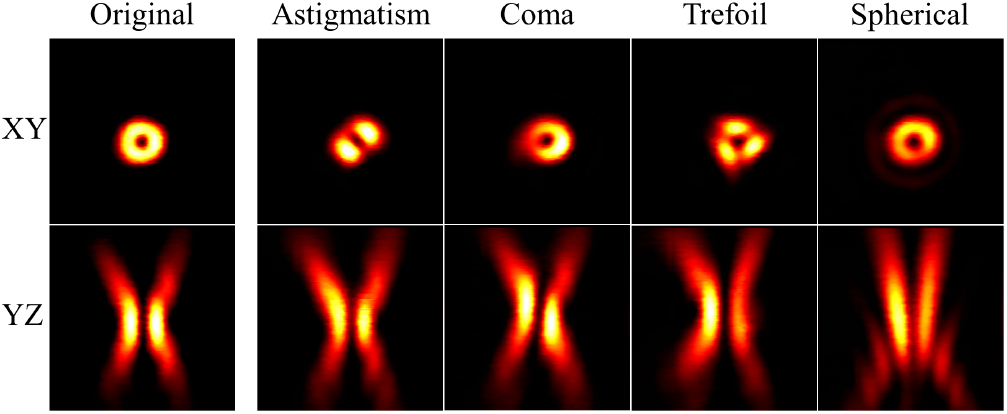
Effect of typical aberrations (astigmatism 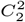, coma 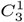, trefoil 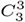 and spherical 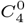) on the donut beam profile. Image size, 3 × 3μ*m*^2^.

While aberrations compromise excitation and STED beams as well as the signal in the emission path, it has been shown that microscope performance is mainly affected by the STED beam, whose central null is particularly sensitive to optical aberrations [10,11,18]. Hence, we decided not to correct excitation and emission wavefronts, but focus on the STED aberrations.

### Acute brain slice preparation

We used the transgenic mouse line *Thy*1 − *H^tg/^*^+^ [26], where a subset of pyramidal neurons in the hippocampus, as well as in cortical layer 4/5, expresses cytosolic YFP. Heterozygous mice were used to obtain a sparser labelling. All procedures were in accordance with the Directive 2010/63/EU of the European Parliament and approved by the Ethics Committee of Bordeaux. Acutely prepared hippocampal slices were obtained from 21-40 day-old mice of both sexes. Mice were anesthetized with isoflurane prior to decapitation and their brains were quickly removed and placed in ice-cold, oxygenated (95% O2 and 5% CO2) NMDG/HEPES-based artificial cerebrospinal fluid (ACSF) containing (in mM): 2.5 KCl, 7.5 *MgSO*_4_, 1.25 *N aH*_2_*PO*_4_, 0.5 *CaCl*_2_, 2.5 *MgCl*_2_, 5 Na-Ascorbate, 3 Na-Pyruvate, 25 glucose, 30 *NaHCO*_3_, 93 NMDG (N-Methyl-D-Glucamine), 93 *HCl* and 20 HEPES (pH 7.4, osmolarity 315 mOsm/L). Transverse 300 μm-thick slices were cut using a vibratome (VT1200, Leica) and incubated for 15 minutes at 33°C in NMDG/HEPES-based solution. Subsequently, slices were transferred into normal ACSF containing (in mM) 125 NaCl, 3 KCl, 26 *NaHCO*_3_, 1.25 *N aH*_2_*PO*_4_, 10 glucose, 2 *CaCl*^2^ and 1 *MgCl*_2_ (pH 7.4, osmolarity 305 mOsm/L), bubbled with carbogen, and allowed to recover at room temperature for at least 1 hour. Slices were maintained and used for a maximum of 4 h after preparation. They were placed on a coverslip with an electrophysiology harp placed on top. The harp was weakly glued to the coverslip with an inert bio-compatible silicon (E43) so that the coverslip could be flipped to be imaged in an upright configuration. The flipped coverslip was then transferred over a recording chamber filled and continuously perfused (2 mL/min) with carbogenated ACSF at room temperature.

### Image processing and analysis

All images were acquired with a pixel size of 19.5 nm (512 × 512 pixels, 10 × 10μ*m*^2^) and a pixel dwell time of 20 *μ*s, which amounts to about 5 s acquisition time per image. Imaging depth into the slice was set by the piezo z-focus, where the fluorescence signal on top of the slice defined the zero level. Image analysis was done on raw data using ImageJ [27]. Images presented in the figures were filtered by a 1-pixel median filter to reduce noise. We measured 2-pixel line profiles a cross t he beads and fitted them with a Lorentzian function, whose full-width at half maximum (FWHM) served as a measure of the spatial resolution. For morphometric analysis of dendritic spines, we used the ImageJ plugin SpineJ, which is based on wavelet filtering and skeletonisation and designed for analyzing super-resolution images of dendritic spines [28].

## Results and discussion

### Calibration of spherical aberration in depth

Using phantom samples, we quantified the spherical aberration as a function of depth. To that end, we used the specific distortions of the donut and bottle beams, observable on the image of gold beads in the agarose gel, as a readout and manually adjusted the Zernike coefficients to retrieve high-quality donut and bottle images. Note that this approach is particularly useful and warranted for complex shaped beams, which are much more sensitive to the effects of aberration than simple Gaussian beams. Fig. 3a displays the schematic of our experiments. The STED beam profile was first optimized, similarly as in Fig. 1b, looking at gold beads placed at the surface of the gel. This allowed correcting for optical aberrations due to imperfections of the microscope. Fig. 3b highlights the distortions of the STED beam profile a t 80 μ m depth without correction (top panel). At this depth, the donut profile was similar to the one observed in the left panel of Fig. 2, especially the side lobes (marked by white arrows), indicating that the main aberration is spherical. Therefore, adjusting the phase mask on the SLM by changing only the spherical Zernike coefficient 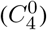 could recover a high-quality STED profile (bottom panel), particularly improving the null (blue arrows). This procedure allowed us to determine the amount of spherical aberration as a function of depth in the phantom sample (Fig. 3c), which exhibits an almost perfect linear behavior as expected for a homogeneous medium [29,30]. Note that beyond this linear dependancy, this approach allows to extract the actual experimental parameters that allows to properly estimate the aberration at a specific depth.

**Fig. 3.**
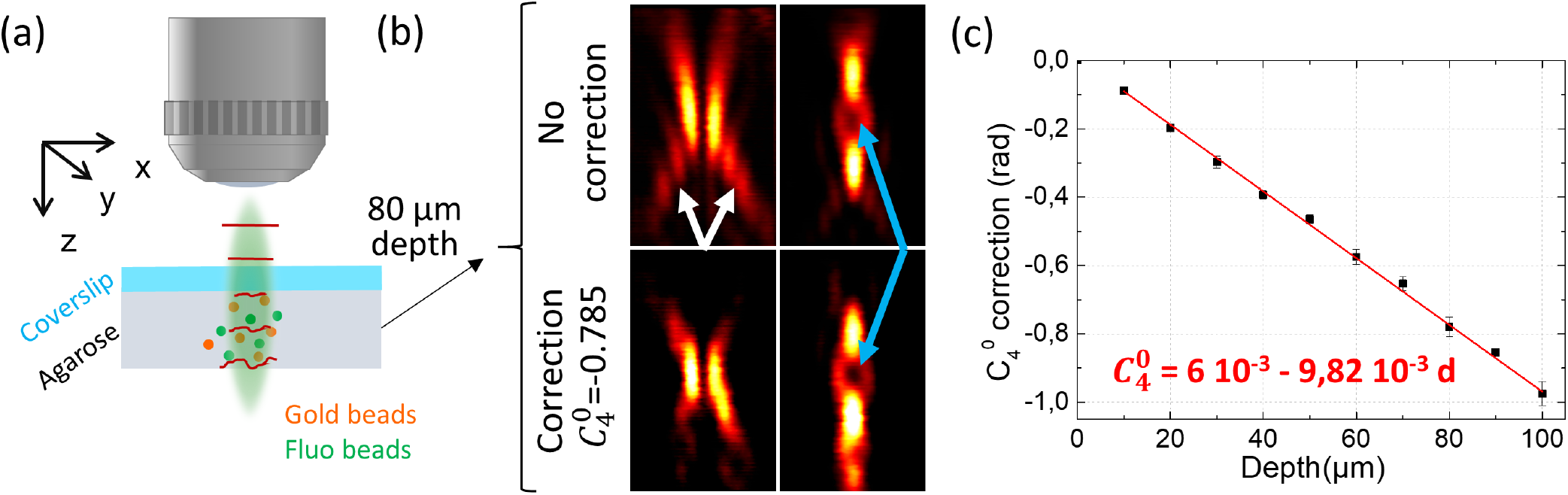
a) Schematic of the phantom sample imaging. b) Donut (left panels) and bottle (right panels) beams side view, visualized using gold beads, at 80 μm depth without and with correction. Image size: 3 × 5μ*m*^2^. (c) Calibration of the spherical aberration as a function of depth in the phantom sample. Averaged from 5 different gels.

It should be noted that this correction is not sufficient to perfectly reshape the STED beam. For example, in Fig. 3b, after correction the donut still exhibits a small distortion typically associated with coma. While we could use a similar approach to quantify coma and astigmatism as a function of depth, these aberrations are more sensitive to sample place ment and thus less reproducible. Nevertheless, since spherical aberration is by far the dominant one, our working hypothesis was that its correction would lead to a significant increase in image quality.

### Resolution improvement in depth

Having determined an effective lateral STED resolution of 68±9 nm using 40 nm fluorescent beads immobilized on a glass slide, we move onto 170 nm diameter fluorescent beads embedded in the agarose phantom sample. While these beads are too large to measure the actual lateral resolution of the STED, we used them because they are very bright and photo-stable, which allowed us to perform reliable repeated measurements. This makes them particularly suitable for the z-scan used to determine the axial resolution. Fig. 4a and b display lateral and axial views of the PSFs measured both in 2P and STED mode at the surface and in depth, illustrating the gain in resolution. While the quality of the STED image appears strongly decreased at a depth of 80 μm, applying the calibrated correction of the spherical aberration allowed us to significantly improve image brightness. This is depicted in Fig. 4c and d, showing the depth dependence of the lateral and axial resolution, respectively. While the effective STED PSF quickly becomes wider in depth (red circles), the resolution remains essentially constant after correction (blue triangles). Notably, the z resolution is more affected than the lateral one, which results from the fact that the bottle beam is more sensitive to aberrations than the donut beam. In contrast, the 2P resolution (black squares) is almost constant, as expected, since the depth range we probed is relatively modest for 2P microscopy. Fig. 4e displays the evolution of the emitted signal as a function of depth and shows that the correction of spherical aberration leads to an increase of 63% in signal strength at 80 μm. Since the 2P intensity is almost constant in this depth range, the loss in signal is mainly due to the decrease in the STED null quality, spuriously de-exciting the fluorescence. It is worth noting that the error bars of the uncorrected STED resolution in Fig. c and d quickly increase in depth. This is due to the strong decrease in signal as illustrated in Fig. 4e, resulting in a poor signal-to-noise ratio. Finally, after correction the bead seems shifted up, as can been seen in Fig. 4b. This is due to the fact that we only corrected for the spherical aberration and not the axial shift introduced by the change in refractive index. Instead, we used the SLM placed in the 2P excitation path (Fig. 1a) to slightly defocus the excitation beam to make it overlap with the STED focus.

**Fig. 4.**
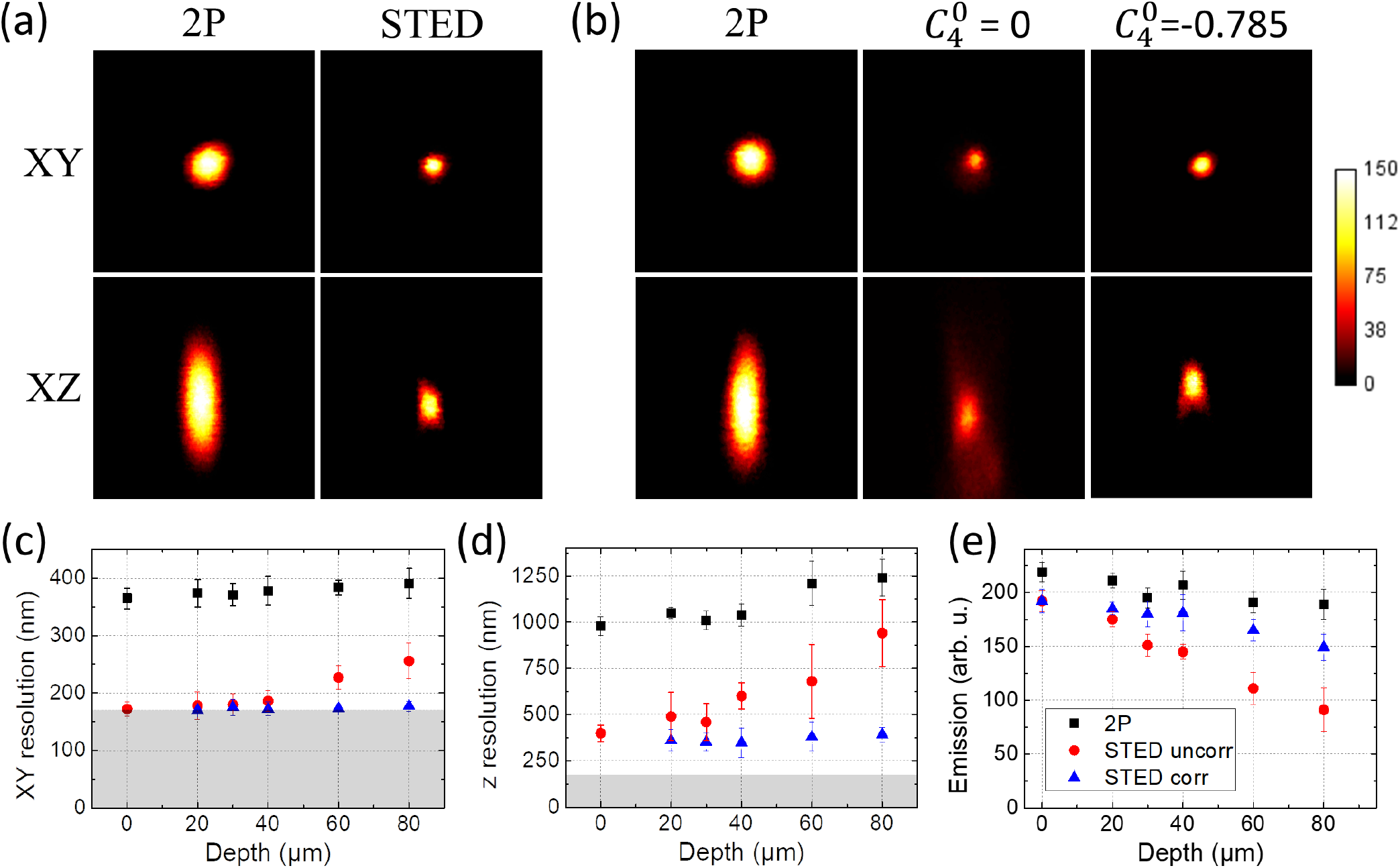
Lateral and axial view of the effective PSFs measured on 170 nm fluorescent beads at the surface of the ghost sample a) and at 80 μm depth b). Image size, 3 × 3*μm*^2^. Evolution of the lateral resolution c), axial resolution, measured as the FWHM, d) and maximum emission signal e) with depth. Averaged from 5 beads at each depth from 3 different gels. Grey zone in c) and d) indicates the inaccessible region of the curve due to the actual beads size.

### Brain slice imaging

Having calibrated the spherical aberration in depth in phantom samples, we investigated if this could be translated into the context of thick biological sample preparations. To that end, we imaged YFP-labelled pyramidal neurons in acute hippocampal mouse brain slices, focusing on dendritic spines. These fine protrusions in the post-synaptic membrane of neurons mediate the vast majority of excitatory synaptic transmission in the brain and their structural and functional plasticity is an important substrate of information processing that underlie sensory perception, motor behaviour and memory, while spine dysfunction is closely linked to brain disorders, such as autism and Alzheimer’s disease [31]. Spines have a characteristic anatomical structure, typically featuring a bulbous head attached to the dendrite via an elongated neck, whose diameter ranges well below 200 nm [32,33]. As a consequence, resolving spine necks requires the use of super-resolution microscopy [34], and imaging them in depth in live settings remains an important challenge [4,35]. Fig. 5a shows images of dendritic spines at different depths acquired in 2P, uncorrected and corrected STED mode, illustrating the gain both in resolution and signal strength obtained with the correction. As all structures on the urface of acute slices inevitably get cut off by the slicing procedure, it is next to impossible to find healthy dendrites within the first ≈ 20*μ*m of the slice. The cellular debris also makes it hard to define the zero level of the tissue surface. To account for this uncertainty, we binned the imaging data into 10 *μ* depth bins.

**Fig. 5.**
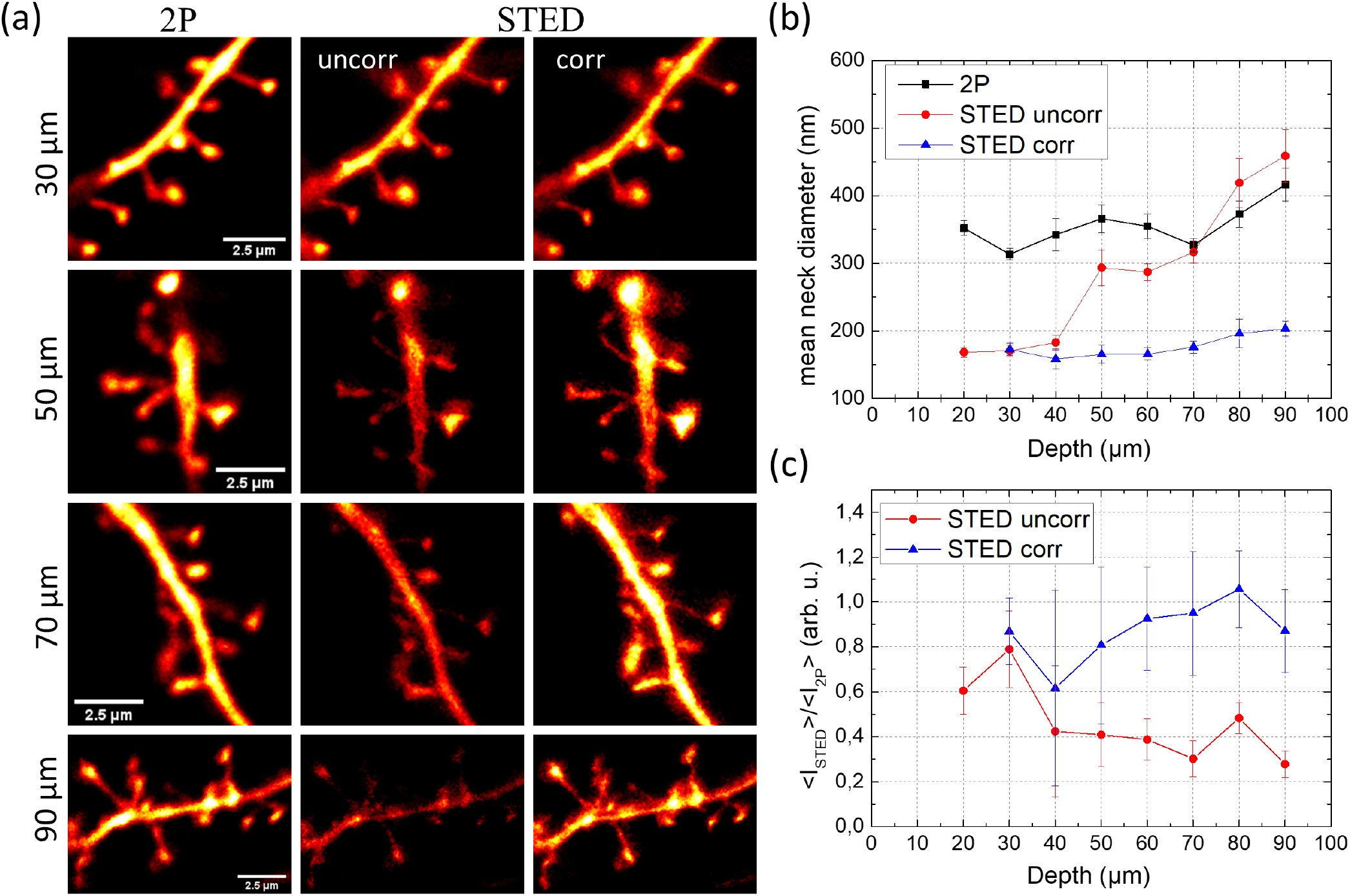
a) Images of dendritic segments of YFP-labelled pyramidal neurons in hippocampal acute slices at various depth. b) Average spine neck diameter measured in 2P, uncorrected and corrected STED mode. c) Average STED intensity detected on the spine heads normalized to the 2P intensity. Measurements were done on 172 spines from 4 different acute slices.

We quantified signal intensity as well as morphological parameters of dendritic spines using SpineJ [28]. Fig. 5b shows that while the uncorrected STED resolution quickly decreased with depth, becoming even worse than 2P beyond 70 *μ*m, the corrected STED allowed us to measure an average spine neck width of 160±15 nm, consistent with what has been reported in the literature [32,34,36] and almost invariant in this depth range. In addition, the relatively small error bars indicates the reproducibility of the measurement across the different brain slices despite there variability. This highlights the fact that, while not offering optimal aberration correction, mimicking brain optical properties as a bulk medium is sufficient to significantly improve the STED resolution (Fig. 5d). The smallest neck widths measurable at 90 μm depth was 81±4 nm, well below the diffraction limit, giving an upper bound of the effective STED resolution at this depth after correction. Note that it was impossible to measure the same neck in the STED image acquired without correction due to poor signal-to-noise ratio, which illustrates the difficulty associated with iterative approach. Also, other morphological parameters, such as head area, spine length and neck length, which are nominally larger than the resolution, were not significantly different from 2P, uncorrected and corrected STED (data not shown). In parallel, Fig. 5c displays the comparison of the corrected and uncorrected STED signal normalized to the 2P intensity. It shows that while the uncorrected STED signal was reduced up to 75% at 90μm depth, the corrected STED image appeared only slightly dimmer (up to 20%) than the 2P image.

Since the mean free path in brain tissue is for photons at 600 nm, i.e. close to our STED wavelength [37,38], after this depth a significant fraction of scattered photons tends to fill up the null of the donut leading to an inevitable decrease of STED performance.

## Conclusion

We have shown, using a phantom sample with adjusted refractive index, that it is possible to calibrate and compensate the distortions of the STED beam profile introduced by spherical aberrations with increasing depth. We then demonstrated that this calibration can be translated into the more complex environment of a living biological sample without any further iterations. This simple approach provides a significant improvement in image quality in depth, offering nanoscale resolution in 3D up to 90 *μ*m inside acute brain slices. Importantly, this approach is not limited to brain samples but could be adapted to other tissues with known and relatively homogeneous refractive indices, and could be applied to other preparations, even potentially in vivo, where approaches based on optical clearing to increase depth penetration cannot be used.

## ACKNOWLEDGEMENTS

This project has received funding from the European Union’s Horizon 2020 research and innovation programme under the Marie Sklodowska-Curie grant agreement No 794492, ANR (ANR-17-CE37-0011), Era-net NEURON (ANR-17-NEU3-0005-01) and FRM (DEQ20160334901). We thank Misa Arizono for comments on the manuscript

